# A novel anti-PD-L1/IL-15 immunocytokine overcomes resistance to PD-L1 blockade and elicits potent antitumor immunity

**DOI:** 10.1101/2022.03.15.484441

**Authors:** Wenqiang Shi, Liangyin Lv, Nan Liu, Hui Wang, Yang Wang, Wen Zhu, Zexin Liu, Jianwei Zhu, Huili Lu

## Abstract

Despite the demonstrated immense potential of immune checkpoint inhibitors in various types of cancers, only a minority of patients respond to these therapies. Immunocytokines designed to deliver an immune-activating cytokine directly to the immunosuppressive tumor microenvironment (TME) and block the immune checkpoint simultaneously may provide a strategic advantage over the combination of two single agents. To increase response rate to checkpoint blockade, in this study we developed a novel immunocytokine (LH01) composed of the antibody against programmed death-ligand 1 (PD-L1) fused to IL-15 receptor alpha-sushi domain/IL-15 complex. We demonstrate that LH01 efficiently binds mouse or human PD-L1 and maintains IL-15 stimulatory activity. In syngeneic mouse models, LH01 showed improved antitumor efficacy and safety versus anti-PD-L1 plus LH02 (Fc-Sushi-IL15) combination and overcame resistance to anti-PD-L1 treatment. Mechanistically, the dual anti-immunosuppressive function of LH01 led to activation of both the innate and adaptive immune response and decreased levels of transforming growth factor-β1 (TGF-β1) within the TME. Furthermore, combination therapy with LH01 and bevacizumab exerts synergistic antitumor effects in HT29 colorectal xenograft model. Collectively, our results provide supporting evidence that fusion of anti-PD-L1 and IL-15 might be a potent strategy to treat patients with cold tumors or resistance to checkpoint blockade.

## 1. Introduction

Therapeutic antibodies that block programmed death-1 (PD-1)/programmed death-ligand 1 (PD-L1) pathway demonstrate cure-like benefits in patients with various types of cancers, but a large proportion of patients experienced a low response rate or rapidly developed resistance to these therapies with relapsed disease ^[1]^. One of the main reasons may be the existence of an immunosuppressive tumor microenvironment (TME), which is caused by altering the immune checkpoint molecule expression, immunosuppressive cytokine secretion, oxygen nutrition status, etc ^[2]^. Cytokines play an indispensable role in regulating immune response, including innate and adaptive immunity, and are the cornerstone of cancer immunotherapy. A variety of immune activating cytokines such as IL-15 have potent anti-tumor efficacy and can markedly prolong the survival periods of patients, which can be combined practically with immune checkpoint inhibitors (ICIs) to address the issue of resistance and increase response rate ^[3, 4]^.

Recombinant human IL-15 was at the top of the National Cancer Institute’s list of potential biopharmaceuticals for tumor immunotherapy in 2008 ^[5]^. IL-15 has a unique mechanism of action in which it binds to IL-15Rα expressed by antigen-presenting cells, then the IL-15/IL-15Rα complex is trans-presented to neighboring NK or CD8^+^ T cells expressing only the IL-15Rβ/γ receptor ^[6]^. In addition to inhibiting IL-2-induced activation-induced cell death, a process that leads to the elimination of stimulated T cells and induction of T-cell tolerance, IL-15 can support long lasting CD8^+^ T cell memory and effector responses against diseased cells ^[7, 8]^. Recombinant IL-15 has demonstrated clinical activity in the treatment of certain cancers, including advanced renal cell carcinoma and metastatic melanoma, and significant increases in the number of memory CD8^+^ T and NK cells were observed in patients’ peripheral blood ^[9, 10]^. There is evidence that increased PD-L1 expression in tumors and decreased IL-15 levels in the TME are correlating with poor clinical outcomes ^[11, 12]^. A clinical trial showed that an IL-15 superagonist, ALT-803, can re-induce immunotherapy response in PD-1-relapsed and refractory non-small cell lung cancer (NSCLC) ^[13]^. Unfortunately, the short half-life and the systemic toxicities of high-dose administration, which can cause fever, fall of blood pressure, flu-like symptoms due to lack of target activity, restrict the further clinical applications of IL-15^[14]^.

Prolonging half-life and increasing targeting ability at the tumor site of this pro-inflammatory cytokine are feasible solutions to the above problems. It has been reported that complexation with the IL-15Rα-sushi domain can improve IL-15 half-life and bioavailability *in vivo* and is effective in mimicking IL-15 trans-presentation ^[15, 16]^. Additionally, the IL-15Rα-sushi domain is a selective and potent agonist of IL-15 action through IL-2/15Rβγ ^[17]^. Immunocytokines are also known as antibody-cytokine fusion proteins, which can utilize the targeting ability of antibody to enrich cytokines at the tumor site. On the one hand, it can enhance tumor targeting capability and reduce the side-effects of cytokines caused by systemic administration. On the other hand, this allows antibodies and cytokines to generate synergistic antitumor effects ^[18, 19]^. Hence, it is a practical strategy to generate an immunocytokine composed of anti-PD-L1 and the IL-15Rα-sushi domain/IL-15 complex to enhance antitumor activity.

In this study, we characterized the biochemical activity of LH01, a bifunctional fusion protein designed to overcome resistance to PD-1/PD-L1 blockade via improving the target activity of IL-15 and blocking the PD-L1 pathway concurrently. The anti-PD-L1 moiety of LH01 is based on atezolizumab, which has been approved to treat different types of cancers ^[20–22]^. We compared the antitumor efficacy of LH01 versus anti-PD-L1+LH02 in murine carcinoma models and preliminarily investigated the mechanism that LH01 overcame resistance to anti-PD-L1 treatment. For the first time, we evaluated the synergistic antitumor effect of combinative administration of LH01 and bevacizumab.

## Results

### Biochemical characterization of the bifunctional fusion protein: LH01

The fusion of IL15Rα-sushi domain (Ile 31 to Val 115) and human IL15 mutant (IL-15N72D) to the C-terminus of the anti-PD-L1 monoclonal antibody was expected to improve the target activity and reduce untoward effects of IL-15 (Figure 1A; Figure S1). A new molecular called Fc-Sushi-IL15 (LH02) was also designed as a non-targeting control (Figure 1A). The mature LH01 protein, whose light chain and heavy chain migrate as approximately 25 kDa and 70 kDa proteins under reducing conditions on SDS-PAGE respectively (Figure 1B), was purified by one-step protein A affinity chromatography (Figure 1C). The purification process of secreted anti-PD-L1 and LH02 protein are the same as that of LH01. Light and heavy chain of anti-PD-L1 migrate as an approximately 25 kDa and 50 kDa protein, and LH02 migrates as an approximately 50 kDa protein under reducing conditions on SDS-PAGE (Figure S2).

**Figure 1.**
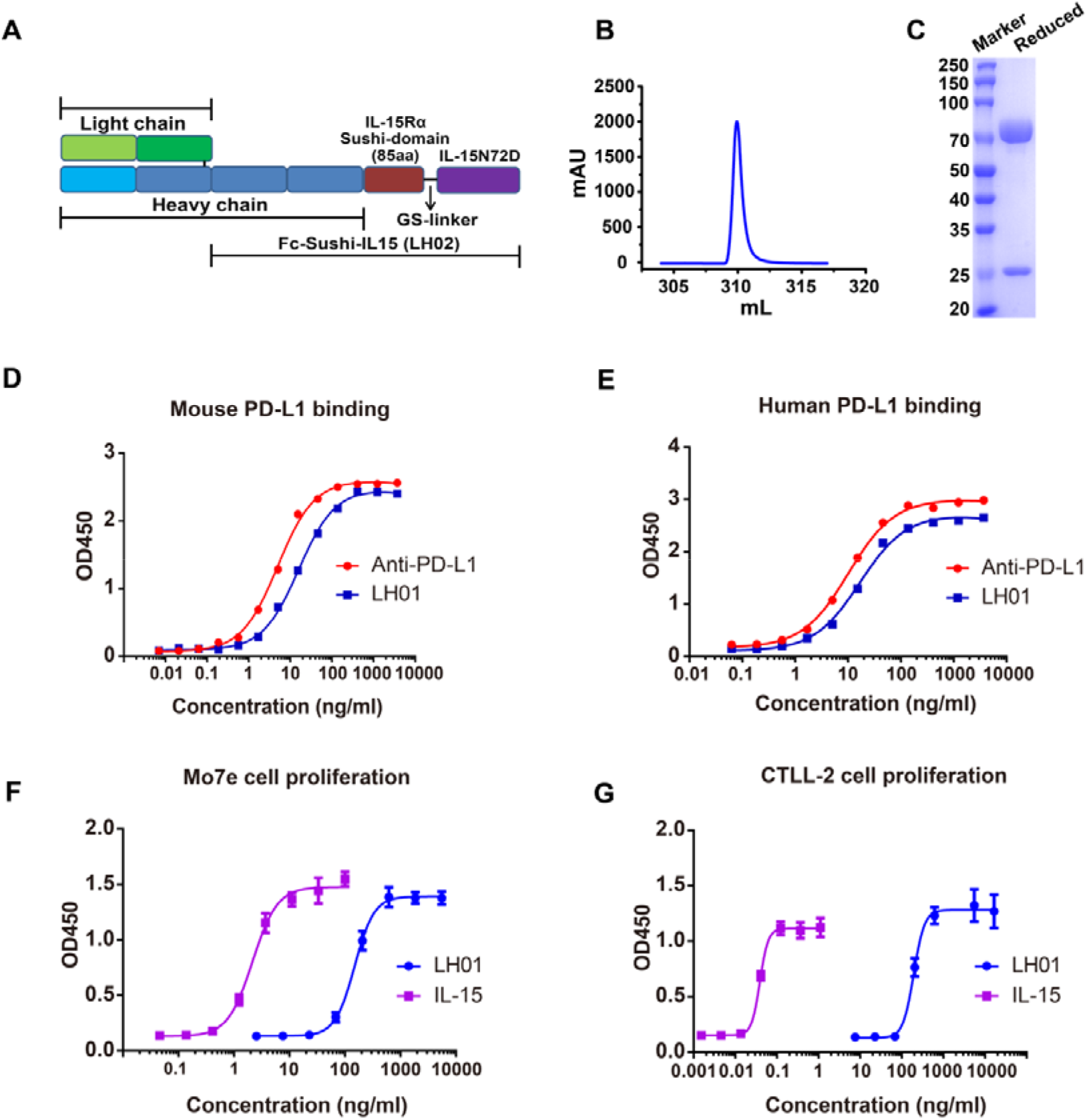
Design, preparation and biochemical characterization of LH01. (A) Schematic representation of fusion proteins: IL-15Rα sushi-domain/IL-15 was directly fused to the C terminus of the Atezolizumab (LH01) or Atezolizumab’s Fc portion (LH02). (B) LH01 was purified by protein A affinity chromatography. (C) Purified LH01 was characterized by reduced SDS-PAGE. (D and E) Binding of LH01 to plate-bound human or mouse PD-L1. Data was analyzed using the one site-total to calculate the EC50 values. (F and G) The biological activity was compared to IL-15 monomer at different concentrations by determining the proliferative potential in human Mo7e cells and murine CTLL-2 cells. Data was analyzed using the four parameter fit logistic equation to calculate the EC_50_ values, and graphs were shown as mean ± SD.

In ELISAs, LH01 bound human or mouse PD-L1 with a profile similar to that of the anti-PD-L1 antibody (EC_50_ = 16.8 and 10.2 ng/mL (or 84.1 and 70.5 pM), 15.9 and 5.0 ng/mL (79.5 and 34.7 pM), respectively), indicating that the binding of the anti-PD-L1 moiety was not affected (Figure 1D and 1E). As shown in Figure 1F and 1G, LH01 exhibited weaker proliferative capacity than IL-15 in human Mo7e cells (EC50 = 149.5 and 2.2 ng/mL (or 0.74 and 0.17 nM)), whereas markedly reduced proliferative activities of LH01 were observed compared to IL-15 in mouse CTLL-2 cells (EC_50_ = 194.5 and 0.039 ng/mL (or 970.4 and 3.03 pM)), which may be explained by the finding that the IL-15Rα-Sushi domain was able to bind IL-15 with high affinity and inhibited proliferation driven through the high affinity IL-15Rα/β/γ signaling complex of the CTLL-2 cells. In a word, LH01 retained strong proliferative capacity in both human Mo7e and mouse CTLL-2 cells.

### Prolonged half-life and improved tumor-targeting distribution of LH01

In order to provide medication guidance for the following animal experiments, we explored the pharmacokinetic properties of LH01. Plasma concentrations of LH01 and IL-15 climbed to peaks and then decreased over time, but that of LH01 decreased markedly slower than IL-15 (Figure 2A). LH01 peaked around 8 h at a concentration of 3231 ng/mL, whereas IL-15 peaked about 1h at a concentration of 39 ng/mL (Table. 1). The half-lives were calculated to be about 12.52 h for LH01 and 1.02 h for IL-15 monomer, indicating that the fusion of IL15 and the sushi domain of the IL15 receptor-alpha to the C-terminus of the anti-PD-L1 monoclonal antibody have markedly prolonged the half-life of IL-15 by more than 12 folds (Table. 1). To further trace LH01 distribution, we collected various tissues 24 h after mice received treatment of LH01. The *in vivo* biodistribution of LH01 displayed a certain specificity, and the concentration of LH01 in tumor tissues was 2 folds higher than that in normal tissues (Figure 2B).

**Figure 2.**
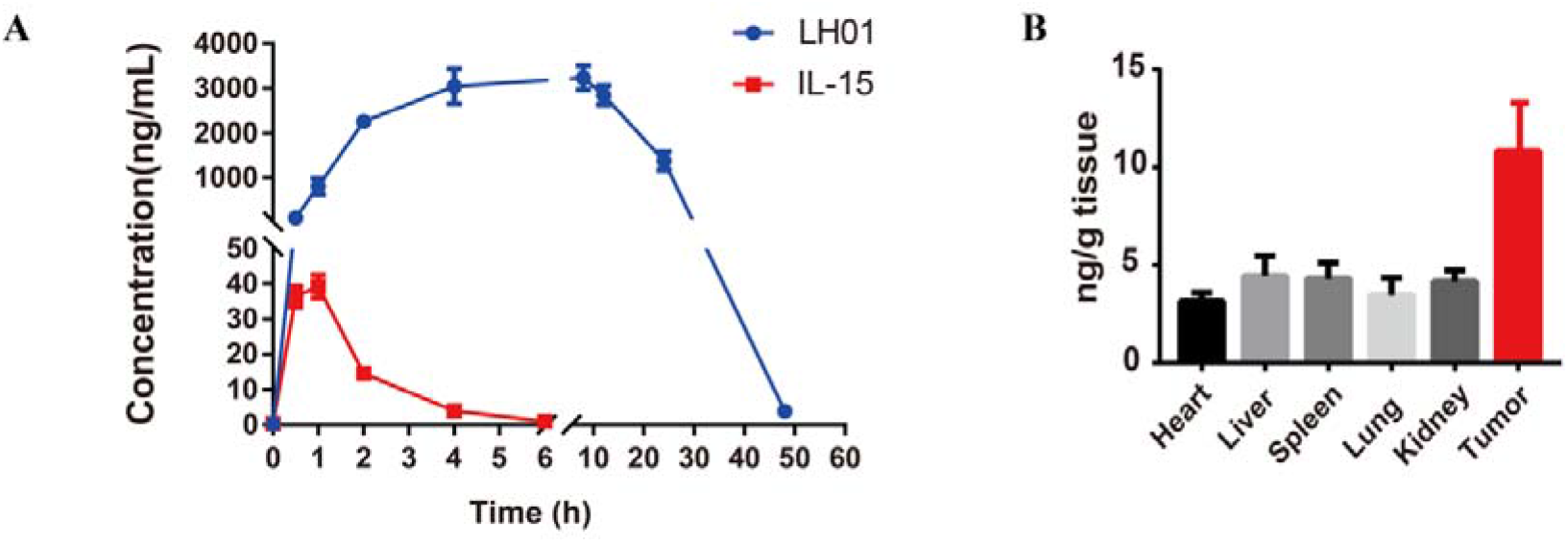
Prolonged half-life and improved tumor-targeting distribution of LH01. (A) Male Balb/c mice aged 9 weeks were intraperitoneally injected with 24.0 μg of LH01 or 3.6 μg of IL-15 (equimolar of IL-15 molecules). The pharmacokinetics curves of LH01 and IL-15 monomer were plotted (n = 5). (B) MC38 tumor-bearing mice (n = 4) were i.p. injected with LH01 at a dose of 1 mg/kg. Tissues were collected at 24 h after injection. The concentrations of LH01 were measured by ELISA. Both graphs show mean±SEM

**Table 1.**
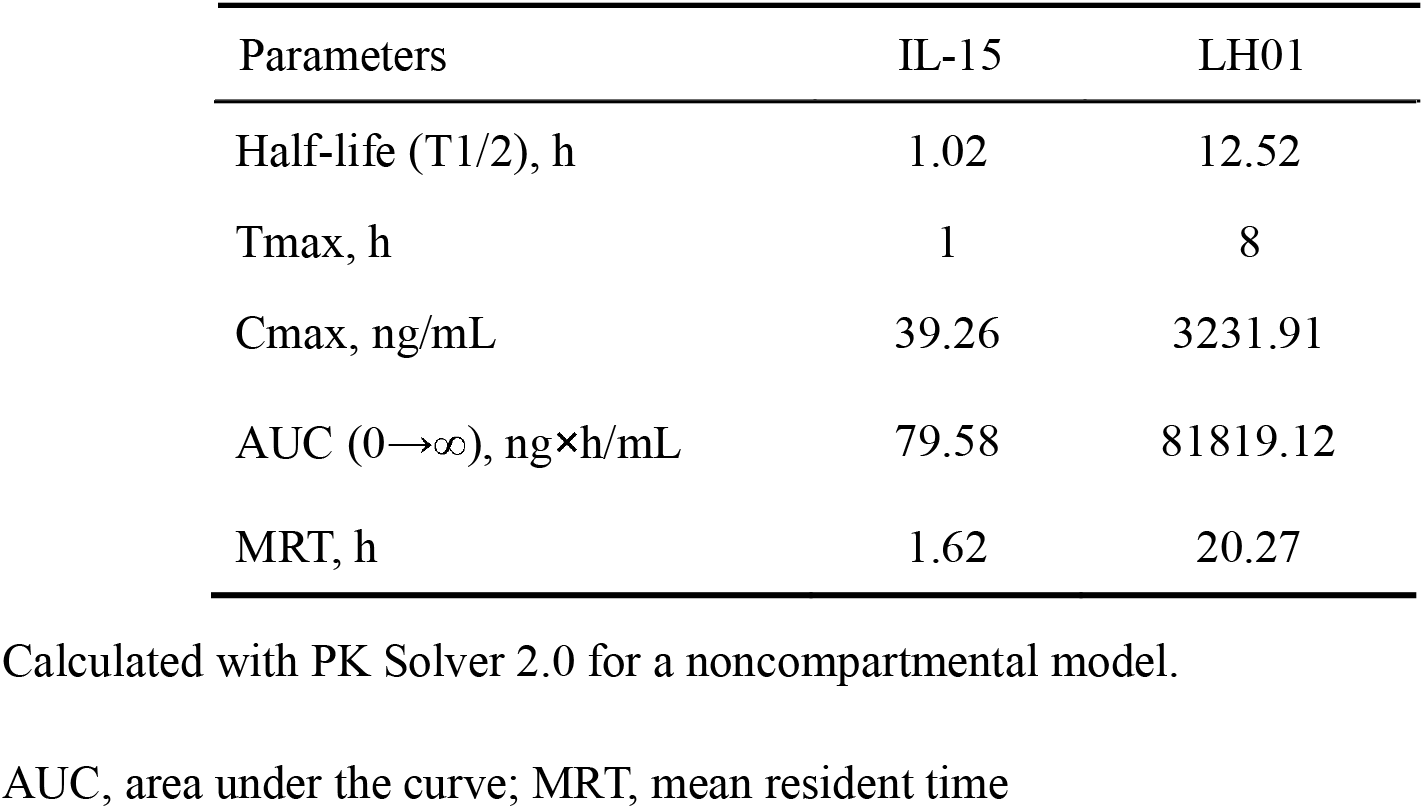
Pharmacokinetic parameters of IL-15 and LH01.

### LH01 improves antitumor efficacy and safety versus anti-PD-L1+LH02 combination

We explored the antitumor effects of LH01 among different doses and meanwhile, we compared the antitumor efficacy of LH01 versus anti-PD-L1 plus LH02 in murine CT26 and MC38 tumor models. Considering that 5 of 8 CT26 tumor-bearing mice died at day 9 after receiving two intraperitoneal treatments of LH02 (1 mg/kg), we decided against using equimolar doses of LH01 and anti-PD-L1+LH02. Instead, LH01 was compared with anti-PD-L1 (5 mg/kg) and LH02 (0.5 mg/kg). LH01 at 3 mg/kg induced similar reduction in CT26 tumor burden compared with 5 mg/kg (TGI: 61.6% (1 mg/kg), 75.3% (3 mg/kg), 79.4% (5 mg/kg)) (Figure 3A), and showed greater decrease in tumor weight than anti-PD-L1+LH02 (Figure 3C). In MC38 tumor models, the immunocytokine demonstrated the antitumor activity in a dose-dependent manner (TGI: 38.5% (1 mg/kg), 56.4% (3 mg/kg), 71.1% (5 mg/kg)), and LH01 (3 mg/kg) exhibited similar antitumor efficacy versus anti-PD-L1+LH02 (Figure 3B and 3C). Notably, in CT26 tumor-bearing mice, anti-PD-L1+LH02 significantly increased the spleen weight compared to PBS group, while there was no obvious spleen weight gain in LH01 group at dose of 1 mg/kg or 3 mg/kg, indicating that LH01 exerted good tumor targeting capability (Figure 3D). LH01 was well tolerated in both tumor models, as neither CT26 nor MC38 tumor-bearing mice obviously lost weight after treatment (Figure 3E). anti-PD-L1+LH02 showed good tolerability in MC38 tumor models, but in CT26 tumor models 2 of 6 mice died after receiving two intraperitoneal treatments due to systemic toxicity. Collectively, these data illustrate that LH01 has greater antitumor activity than anti-PD-L1+LH02 and a favorable tolerability.

**Figure 3.**
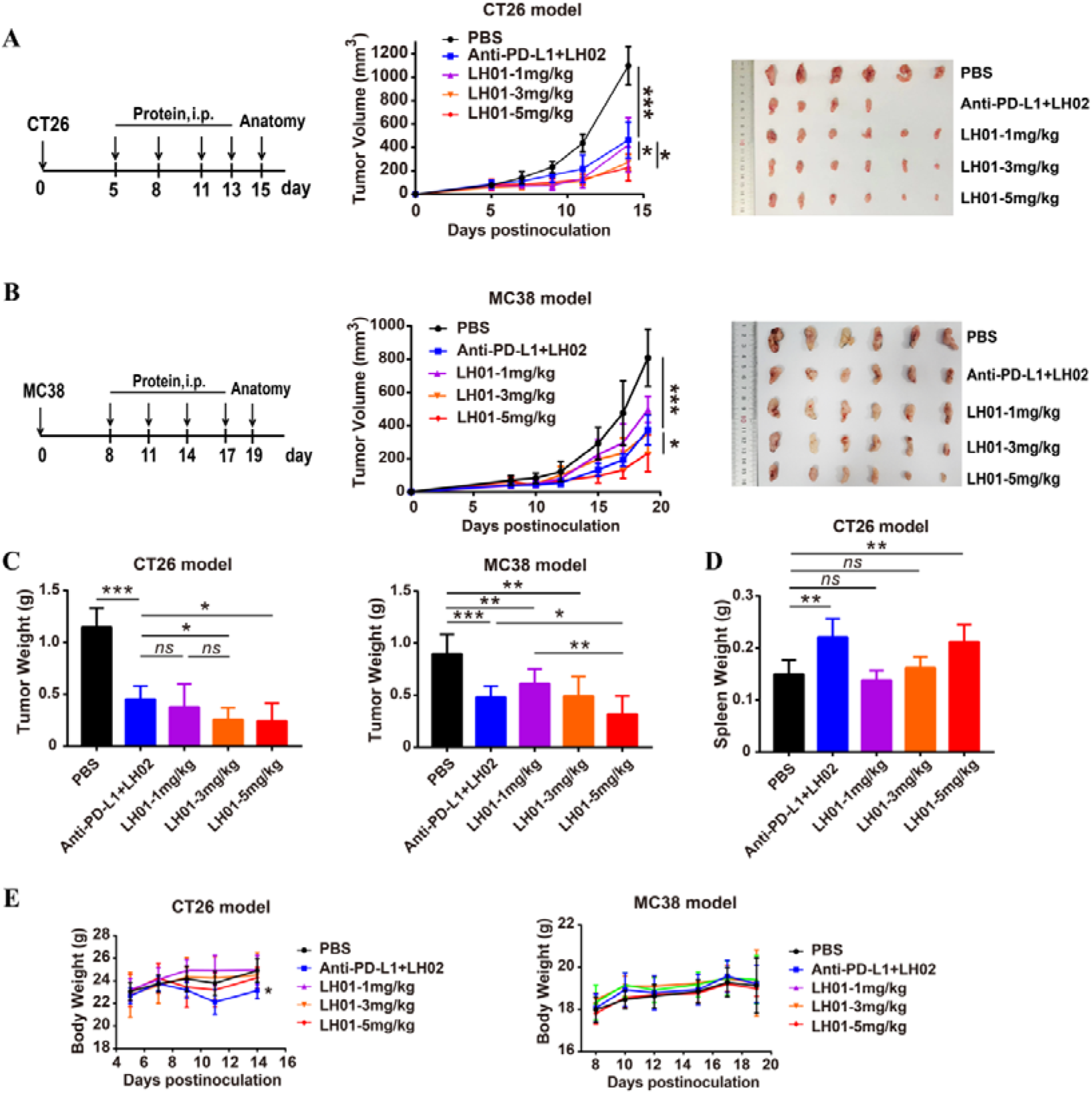
LH01 reduces CT26 and MC38 tumor burden and ameliorates safety versus LH02+anti-PD-L1. (A and B) CT26 tumor cells (1×10^6^, subcutaneously) and MC38 tumor cells (5×10^5^, subcutaneously) were implanted into the right flank of female Balb/c and C57BL/6 mice, respectively. Mice were randomized into 5 groups based on tumor size and treatment initiated when tumors reached 50-100mm^3^. Mice were treated with PBS, LH02+anti-PD-L1 or LH01, and the progression curves of tumor volumes were depicted (n = 6). Tumors were removed and photographed after euthanasia. C, Tumors were weighed. D, Spleens of Balb/c mice were removed and weighed after euthanasia. F, Body weights of mice was recorded. All graphs show mean±SD.

### LH01 induces both innate and adaptive immune cell activation in tumors

IL-15 is a pleiotropic cytokine that plays a vital role in regulating innate and adaptive immunity and can strongly expand CD8^+^ T and NK cells with much weaker regulatory T cells (Tregs)-stimulating activity ^[23]^. To explore the changes in splenic and intratumoral CD8^+^ T, NK and Tregs populations, we performed flow cytometry analysis of dissociated tumors and spleens from CT26 tumor-bearing mice. LH01 markedly increased the CD8^+^ tumor-infiltrating lymphocytes (TILs) compared with PBS or LH02+anti-PD-L1, which suggested that LH01 can selectively activate CD8^+^ T cells in the tumor (Figure 4A). Besides its effects on CD8^+^ T cells, LH01 treatment also increased the tumor-infiltrating natural killer cells (TINKs) and dramatically decreased the tumor-associated Tregs (Figure 4B and 4C). In comparison with PBS group, LH01 treatment only elicited slight increase in splenic CD8^+^ T cells, while anti-PD-L1+LH02 treatment markedly increased splenic CD8^+^ T and NK cells, which indicated good tumor targeting activity of LH01 (Figure 4D, Figure S3). To our surprise, we found that LH01 treatment remarkably increased splenic Tregs versus PBS group, which may be beneficial to reduce systemic toxicity (Figure 4D).

**Figure 4.**
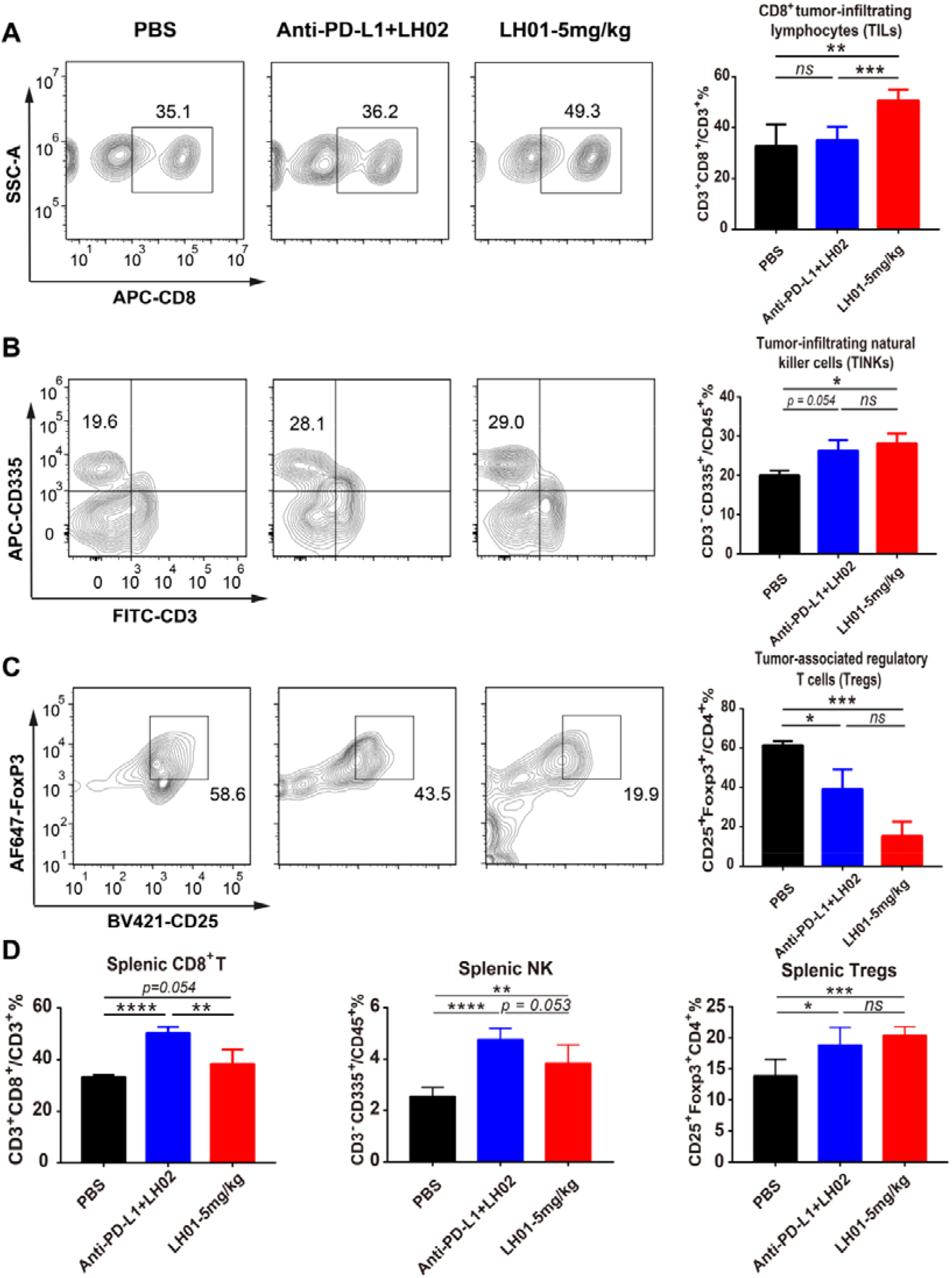
LH01 increases both adaptive and innate immune cell activation in tumors and spleens. Flow cytometry analysis of dissociated tumors and spleens from CT26 tumor-bearing mice treated as described in Figure 3. (A-D) The percentages of intratumor (A-C) and splenic (D) CD8^+^ T cells, NK cells and Tregs were shown for populations of CD3^+^, CD45^+^ and CD4^+^ lymphocytes. Data are reported as the mean ± SEM.

### LH01 overcomes resistance to PD-1/PD-L1 blockade in MC38 model correlated to inhibition of TGF-β1

Our results showed that mice did not respond to anti-PD-L1 treatment at a dose of 10 mg/kg with primary resistance to therapy, while LH01 displayed obvious therapeutic improvements (Figure 5A). Both anti-PD-L1 and LH01 treatments relieved inhibition of T cell via PD-1/PD-L1 axis, and showed a significant increase in CD8^+^ TILs than control group (Figure 5B). The above meant that impairment of T cell function caused by immunosuppressive TME may contribute to resistance to PD-L1 blockade. As an important feature of TME, high reactive oxygen species (ROS) is detrimental to the survival and function of T lymphocytes ^[24]^. We explored whether LH01 can inhibit the apoptosis induced by oxidative stress. The results of flow cytometry demonstrated that the apoptosis rate was significantly higher than that of the control group (57.4 ± 3.0 % VS 23.9 ± 1.7 %) after T lymphocyte cell line CTLL-2 was incubated with 50 μM H_2_O_2_ for 18h, whereas the addition of LH01 could dramatically reverse the apoptosis induced by H_2_O_2_ (57.4 ± 3.0 % VS 6.2 ± 0.6 %) (Figure 5C).

**Figure 5.**
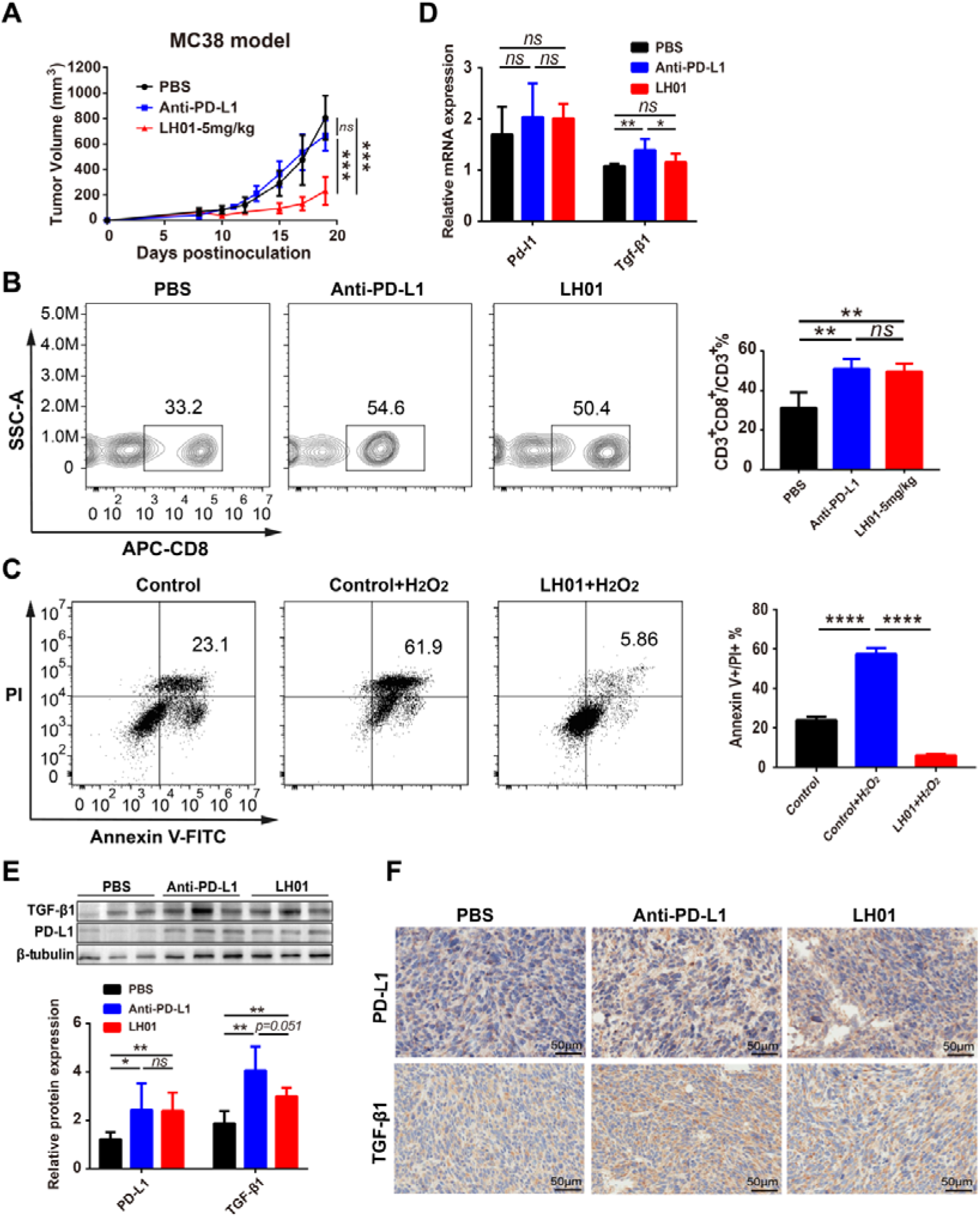
LH01 overcomes resistance to anti-PD-L1 treatment related to suppression of TGF-β1. (A) C57BL/6 mice transplanted s.c. with MC38 cells (right) were treated with PBS, anti-PD-L1 or LH01 as described in Figure 3, and the progression curves of tumor volumes were depicted (n = 6 mice/group). (B) The percentage of CD8^+^ TILs was shown for populations of CD3^+^ lymphocytes. (C) Well-grown CTLL-2 cells were planked with 2*10^5 cells per well on 12-well cell culture plates. Flow cytometric analysis of CTLL-2 cells untreated (left), treated with 50 μM H_2_O_2_ (middle) or 50 μM H_2_O_2_ + LH01 (right) for 18 hours. The proportion of late apoptotic cells is statistically analyzed (n = 4). (D and E) Quantitative real-time PCR analysis and western blot were performed to measure the expression levels of PD-L1 and TGF-β1 in tumor tissues. (F) tumor tissues were fixed, followed by immunohistochemical staining for PD-L1 and TGF-β1. Data are reported as the mean ± SD.

Transforming growth factor-β (TGF-β) exerts diverse effects in tumorigenesis and progression. The pleiotropic nature of TGF-β signaling within the TME facilitates tumor immune escape and promotes tumor progression via induction of epithelial-mesenchymal transition, angiogenesis, and stromal modification ^[25, 26]^. It has revealed that TGF-β participates in the resistance to PD-1/PD-L1 blockade ^[27, 28]^. Anti-PD-L1 treatment markedly increased expression levels of PD-L1 and TGF-β1, which partly explained the resistance to its treatment (Figure 5D-5F). Intriguingly, compared to anti-PD-L1 group, LH01 treatment did not significantly alter PD-L1 expression levels, but remarkably reduced TGF-β1 levels (Figure 5D-5F). The results suggested that inhibition of TGF-β signaling may target mechanisms of resistance and sensitize tumors to immunotherapy.

### Combination therapy with LH01 and bevacizumab exerts synergistic antitumor effect

Previous studies have displayed that angiogenesis is an essential process for the proliferation of solid tumor and VEGF can elicit immunosuppressive effects in the TME, suggesting that anti-angiogenic agents and LH01 could generate synergistic antitumor efficacy ^[29, 30]^. Bevacizumab is a molecularly targeted drug that can inhibit tumor angiogenesis by binding to vascular endothelial growth factor A (VEGF-A) around tumor ^[31]^. In a HT29 xenograft model, mice experienced a slight and significant reduction in tumor volume and tumor weight after receiving bevacizumab and LH01, respectively (Figure 6A and 6B). Two mice died after receiving four intraperitoneal treatments of LH01, possibly due to graft-versus-host disease caused by infused human peripheral lymphocytes (Figure 6C). Combination therapy of LH01 and bevacizumab markedly reduced tumor volume and tumor weight (Figure 6A and 6B) versus LH01 or bevacizumab and showed synergistic antitumor activities (CI = 1.09). Larger areas of necrosis were observed in the combination regimen than the other three groups (Figure 6D). Both LH01 monotherapy and combination therapy obviously reduced the expression levels of Ki67 compared to PBS or anti-VEGF treatment, which demonstrated a poor proliferative and metastatic ability of tumor cells (Figure 6D). Our results indicate that LH01 is a promising candidate to exert enhanced antitumor activities in combination with angiogenesis inhibitors.

**Figure 6.**
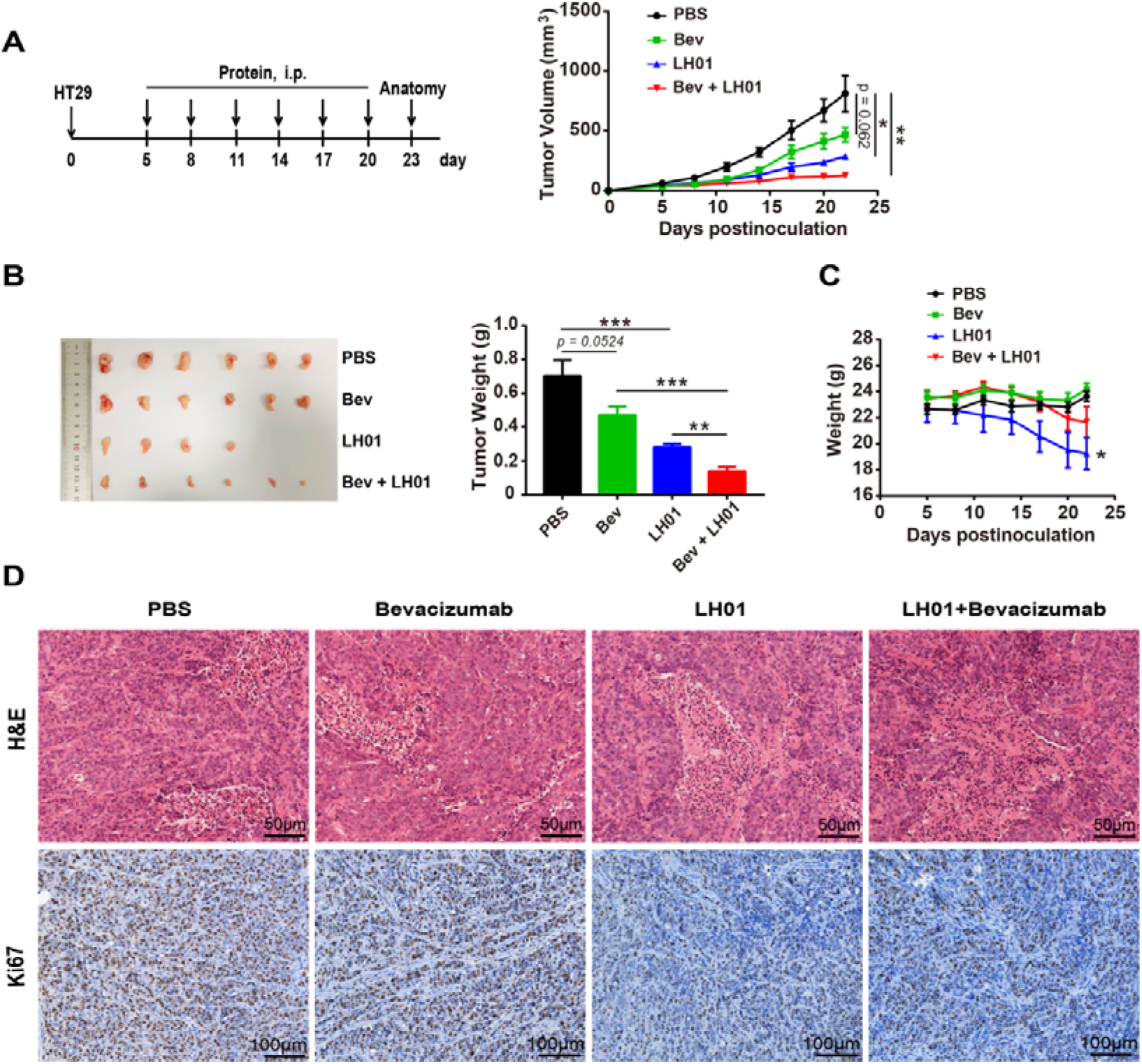
Combining LH01 with bevacizumab enhances antitumor activity. (A and B) NOD-SCID mice were inoculated subcutaneously with 3 × 10^6^ HT29 cells, and subsequently received 3 × 10^6^ fresh human PBMCs intravenously on the same day. Mice were randomized into 4 groups and treatment initiated when tumors reached 40-80mm^3^. Mice were treated intraperitoneally with LH01 (3 mg/kg), bevacizumab (10 mg/kg) or LH01 (3.0 mg/kg) + bevacizumab (10 mg/kg) at days 5, 8, 11, 14, 17 and 20 (n = 6). (A) Tumor volumes were measured every 3 days. (B) Tumor weight (day 23). (C) Body weights of mice. (D) tumor tissues were fixed, followed by H&E staining and immunohistochemical staining for Ki67. CI was calculated based on the formula Ea+b/(Ea + Eb - Ea×Eb). Data are reported as the mean ± SEM.

## Discussion

The antibodies targeting the immune checkpoint have become the protagonist of immunocytokines with the breakthrough progress of immune checkpoint inhibitors for tumor treatment in the past few years. Recently, Martomo et al. preliminarily explored the antitumor activity of anti-PD-L1/IL-15 fusion protein KD033 on various solid tumor models in mice, whereas they did not provide further rationale for KD033 to treat patients with cold tumors or resistance to ICIs ^[32]^. LH01 is different in structure compared to KD033: the antibody part is atezolizumab and the sushi domain is 85 amino acids in length with higher binding affinity to IL-15 than the 65 amino acids of KD033 ^[15]^. Interestingly, our results show that LH01 can overcome primary resistance to PD-1/PD-L1 blockade by down-regulating TGF-β1 levels within the TME without markedly affecting PD-L1 expression. Additionally, we demonstrate that LH01 can induce the development of an inflamed TME through enhancing the populations of CD8^+^ TILs and TINKs with a decrease in Tregs populations. Resistance is a major obstacle to cancer immunotherapy, and its mechanisms are varied and complicated. Amelioration of primary resistance to anti-PD-L1 therapy by using LH01 may be related to converting inherently immunosuppressive TME to immunosupportive one.

The format of immunocytokine has a significant impact on its targeting activity. The homodimeric format usually possesses a high binding avidity to the target and a long residence time at the tumor site. In our study, we found that the LH01 did not substantially increase weight of the spleen of tumor-bearing mice at the dose of 1 mg/kg and 3 mg/kg compared to that of control group, which indicated that the LH01, a homodimeric fusion protein, had good tumor targeting activity. It should be noted that LH01 has a favorable safety at a high dose (5 mg/kg) in both CT26 and MC38 models, with no difference observed in mice body weights. On the other hand, fast blood clearance profiles may be beneficial to reduce untoward effects associated with the use of potent pro-inflammatory cytokine payloads, which perhaps partially explains the good safety of LH01 in mice.

In principle, certain immunocytokine products could mimic the action of bispecific antibodies (BsAbs). The cytokine moiety can engage in a binding interaction with its cognate receptor on the surface of T cells, thus creating an immunological synapse with the tumor cells ^[33]^. Bispecific T cell engagers (BiTEs), one kind of BsAbs, have been attracting a great deal of attention due to its unique mechanism of action and significant antitumor activity. BiTEs demonstrated remarkable efficacy in B cell hematologic malignancies, but the use of such new drugs to treat solid tumors is unsatisfactory ^[34]^. Although BiTEs can redirect T cells to specific tumor antigens and activate T cells directly, the immunosuppressive factors in the TME, including high levels of ROS, hypoxia, TGF-β, etc, are not conducive to the proliferation and survival of T and NK cells in solid tumors, which can importantly reduce its antitumor activity ^[35, 36]^. However, the pro-inflammatory cytokine moiety of immunocytokine can convert the tumor immunosuppressive microenvironment to a certain extent, and promotes the activation and proliferation of T and NK cells, which is supported by our results that LH01 can inhibit the apoptosis of CTLL-2 under high levels of ROS and down-regulate the TGF-β1 levels in TME. In terms of BiTEs, a major restriction of tumor-associated antigen selection in solid tumors is that low-level expression is often found in normal tissue exposing the patients to a risk of “on-target, off-tumor” toxicity ^[36]^. In the case of immunocytokines, it seems not so demanding for antigen specificity, and the adverse effects of immunocytokines are mainly caused by cytokine moiety.

Given that LH01 is well tolerated in preclinical models, we believe that this bifunctional fusion protein represents a promising candidate for inclusion in combination therapy regimens. We have validated this in our murine models, in which combining LH01 with a VEGF-A inhibitor bevacizumab elicits enhanced and superior antitumor activity over that of either agent alone. Vascular abnormalities resulting from elevated levels of proangiogenic factors (e.g. VEGF and angiopoietin 2) are a hallmark of most solid tumor ^[37]^. Additionally, proangiogenic factors have been reported to play a vital role in immunosuppressive TME ^[38]^. For instance, VEGF can directly elevate PD-L1 expression on dendritic cells resulting in impaired function of T cells, and VEGF can also directly binds to VEGFR2 on regulatory T cells (Tregs) and myeloid-derived suppressor cells (MDSCs), which increases these immunosuppressive cells into TME ^[39, 40]^. Our results further indicate that the combination of the other inhibitors of VEGF signaling pathway including small molecule receptor tyrosine kinases inhibitors (sunitinib, sorafenib, and pazopanib) with LH01 has the potential to generate greater antitumor effects.

Our study has some limitations. First of all, the mechanisms that LH01 overcomes resistance to anti-PD-L1 remains to be further studied because a variety of factors including other immune checkpoints, cancer neoantigens, soluble MHC related molecules, and cytokines in the TME also affect anti-cancer immune response ^[41]^. In addition, we noted that the CT26 tumor-bearing mice showed slightly ungroomed hair without weight loss after third administration of LH01 at a dose of 5 mg/kg, which was associated with side effects caused by cytokine IL-15. Given that most of immunocytokines still produce the same adverse effects as cytokines in clinical trials ^[42]^, further efforts should be made to improve safety by structure-based design.

In conclusion, LH01 elicits superior antitumor efficacy and a good safety profile in preclinical models. LH01 possesses the potential to help T cells resist damage from unfavorable factors and overcome primary resistance to PD-1/PD-L1 blockade by inhibiting TGF-β1 within the TME, which offers supporting evidence for clinical use of LH01 for treatment of patients with resistance to ICIs or cold tumors. LH01 can be combined more practically with other therapies to target even more pathways to improve clinical benefit. Altogether, LH01 represents a potential candidate for further clinical investigation.

## Materials and Methods

### Cloning, expression, and purification

The plasmids encoding LH01, LH02, and anti-PD-L1 were constructed as shown in Figure S1. The DNA sequences of IL-15 mutant (IL-15N72D) and IL-15Rα sushi-domain (Ile 31 to Val 115) were amplified by polymerase chain reaction using the pIL-15 and psIL-15Rα/Fc we reported previously as template ^[43]^. All the plasmids were constructed by inserting the DNA fragments into the vector we used before ^[43]^. The light and heavy chain expression plasmids of LH01 or anti-PD-L1 were mixed at 2:1 and co-transfected using liner polyethylenimine (PEI) with a molecular weight of 25 kDa (Polysciences, Warrington, PA, USA). LH02 was produced by transfecting HEK293E cells with Fc-Sushi-IL15-expression plasmid alone. LH01, anti-PD-L1 and LH02 were all purified by affinity chromatography using a protein A affinity column (GE Healthcare, Piscataway, NJ, USA) and analyzed in reducing condition on sodium dodecyl sulfate-polyacrylamide gel electrophoresis (SDS-PAGE).

### Cell lines

HEK293E, CTLL-2 cell lines was kept in our laboratory and cultured as previous descriptions ^[43]^. Mo7e, MC38 and CT26 murine colon carcinoma cell lines were obtained from the American Type Culture Collection (ATCC, Manassas, VA, USA). Mo7e cells were grown in RPMI 1640 (Gibco, Waltham, MA, USA) containing 10% FBS (Gibco) and 10ng/mL human GM-CSF (Sino Biological, Beijing, China). MC38 and CT26 cells were maintained in Dulbecco’s Modified Eagle Medium (DMEM) containing 10% FBS. All cells above were maintained under aseptic conditions and incubated at 37°C with 5% CO_2_.

### Measurement of LH01 binding and pharmacokinetics by enzyme-linked immunosorbent assays (ELISAs)

#### ELISAs for PD-L1 binding

ELISAs were performed using standard methods. Briefly, 96-well ELISA plates (Corning, Corning, NY, USA) were coated by incubating with 1.0 μg/mL of recombinant human or mouse PD-L1 (Novoprotein, Nanjing, China) at 4°C overnight, then washed four times with PBST (PBS, 0.05% Tween-20) and blocked with 5% bovine serum albumin for 2 hours at room temperature. After washing the plates, serial dilutions (1:3) of LH01 and anti-PD-L1 were added to the plates in duplicate and incubated at room temperature for 2 hours. Plates were washed four times and incubated with Peroxidase AffiniPure Goat Anti-Human IgG (H+L) (Jackson ImmunoResearch, West Grove, PA, USA, 1:10,000 dilution) at room temperature for 1 hour. After being washed, TMB single component substrate solution (Solarbio, Beijing, China) was added to the plates and incubated in the dark for 3-5min. After terminating the reaction with 2 M sulfuric acid, absorbance was read at 450 nm.

#### Pharmacokinetic evaluation of LH01and IL-15 by ELISAs

Plasma samples were drawn from mice 0.5, 1, 2, 4, 8, 12, 24, and 48 hours after treatment with LH01, and 0.5, 1, 2, 4, and 6 hours after treatment with IL-15 monomer. A 96-well ELISA plate, previously coated overnight at 4°C with 1.0 μg/mL of recombinant human PD-L1, was incubated with plasma samples for 2 hours from mice treated with LH01. The following experimental procedure was the same as described above. The human IL-15 ELISA Pair Set (Sino Biological) was used for the quantitative determination of IL-15 monomer.

### Cell Proliferation Assay

Mo7e cells were wash with human GM-CSF free medium (RPMI1640 + 10% FBS) and seeded into 96-well plate with 2×10^4^ cells in a volume of 50 μL per well. After 4 hours’ starvation, serial dilutions (1:3) of LH01 or IL-15 was added to the plate in sextuplicate at 50μL per well to achieve a final density of 2×10^4^ cells/100 μL/well. After being incubated for 96 hours at 37°C with 5% CO_2_, the cell viability was measured using Cell Counting Kit-8 (Dojindo, Kumamoto, Japan). Absorbance was read at 450 nm, and the final OD450 value was calculated as the reading of sample well minus the reading of blank well containing medium. The method used in CTLL-2 cell proliferation assay was identical to that of Mo7e, except that the number of cells used was 1×10^4^ and the incubation time was 72 hours.

### Animal experiments

Female Balb/c, C57BL/6 and NOD-SCID mice aged 6-8 weeks were purchased from Shanghai SLAC Laboratory Animal Co., Ltd and reared under specific pathogen-free conditions. All experiments were approved by the Animal Care and Use Committee of Shanghai Jiao Tong University. All mice were treated humanely throughout the experimental period. Human peripheral blood mononuclear cells (PBMCs) were isolated by Ficoll density gradient centrifugation to serve human T lymphocytes with procedures previously we described ^[45]^. For antitumor studies, tumors were measured every two days using digital caliper, and volumes were calculated as (length × width^2^)/2. Tumor Growth Inhibition (TGI): TGI(%) = 100×(1-T/C). T and C were the mean tumor volume of the treated and control groups, respectively.

### Flow cytometric analysis of splenic and intra-tumoral CD8^+^ T, NK and regulatory T cells

150 mg tumor tissues were finely minced and digested with 4 mL lysis solution (2 mg/mL collagenase IV and 1.2 mg/mL hyaluronidase). The digested tumor tissues were filtered through 200-mesh nylon net to obtain the cell suspension. Centrifuge, then discard the supernatant, and wash the cells once with 6 mL FACS buffer (PBS + 2% FBS). The cells were re-suspended in 6 mL FACS buffer, and filtrated through 200-mesh nylon net again to obtain pre-treated single cell suspension. The spleens were gently grinded and lymphocytes were isolated with lymphocyte separation medium (Dakewe, Beijing, China).

Cell samples were blocked with anti-mouse CD16/CD32 mAb 2.4G2 (BD Biosciences, San Jose, CA, USA) at 4°C for 15 min and incubated with surface marker antibodies at 4°C for 25 min. For the detection of Tregs, cell samples would be further incubated with FOXP3 Fix/Perm buffer (BioLegend, San Diego, CA, USA) for 20 min and FOXP3 Perm Buffer for 15 min at room temperature before anti-FOXP3 was added.

The antibodies and reagents were used as follows: anti-mouse CD45-Percp/Cyanine 5.5 (BioLegend), anti-mouse CD45.2-PE (BioLegend), hamster anti-mouse CD3e-FITC (BD Biosciences), rat anti-mouse CD4-PE (BD Biosciences), rat anti-mouse CD8a-APC (BD Biosciences), rat anti-mouse Nkp46-Alexa Flour 647 (BD Biosciences), anti-mouse CD25-Brilliant Violet 421 (BioLegend), anti-mouse/rat/human FOXP3-Alexa Fluor 647 (BioLegend). Flow cytometry was performed on a CytoFLEX cytometer (Beckman Coulter) and analyzed by FlowJo 10 (TreeStar, Ashland, OR, USA).

### Flow cytometric analysis of cell apoptosis

Flow cytometry was performed to detect the apoptosis of CTLL-2 cells by using Annexin V-FITC/PI Apoptosis Detection Kit (Vazyme) and analyzed by FlowJo 10 (TreeStar).

### RNA isolation and qRT-PCR analysis of mRNA expression

Total RNAs of pretreated tumor tissues were extracted by using Ultrapure RNA Kit (Cwbio, Beijing, China). cDNA was synthesized using a PrimeScript RT Master Mix (Takara, Tokyo, Japan), and quantitative real-time polymerase chain reactions (qRT-PCR) were analyzed on an Applied Biosystems 7500 Fast Real-Time PCR System (ThermoFisher Scientific, Eugene, OR, USA) using Hieff^®^ qPCR SYBR^®^ Green Master Mix (Yeasen, Shanghai, China). The primer sequences are listed in table S1. All results were normalized to GAPDH expression and calculated using the 2^-(ΔΔCt)^ method.

### Western blotting

MC38 tumor tissues were lysed using radio immunoprecipitation assay buffer (Beyotime, Shanghai, China). Protein lysates were separated on 10% SDS-PAGE gels and then transferred to PVDF membranes (Millipore, Billerica, MA, USA). The membranes were blocked with 5% nonfat dry milk at room temperature for 2 hours and then incubated at 4°C overnight with primary antibodies against β-tubulin (Abcam, Cambridge, MA, USA), PD-L1 (ABclonal, Wuhan, China) or TGF-β1 (Abcam). Membranes were washed three times and incubated with HRP-conjugated secondary antibodies. Target proteins were visualized using ECL (ThermoFisher Scientific). The autoradiograms were analyzed with Image J software to quantify the band densities.

### Histopathological and IHC analysis

The tumor tissues were fixed in 4% paraformaldehyde, and then embedded in paraffin, sectioned (4 μm), and stained with hematoxylin and eosin. After dewaxing and hydration, the tumor sections were treated with heat induced epitope retrieval and 3% hydrogen peroxide for 15 minutes to block the endogenous peroxidase activity. Next, the tumor sections were blocked with 5% BSA for 30 min and incubated with anti-mouse PD-L1 rabbit antibody (ABclonal), anti-mouse TGF-β1 rabbit antibody (Abcam), or anti-human Ki67 rabbit antibody (Servicebio, Wuhan, China) at 4°C overnight. Afterward, the sections were incubated with the HRP-conjugated goat anti-rabbit secondary antibody (Servicebio) for 50 minutes. Finally, the sections were stained with DAB detection kit (Dako, Copenhagen, Denmark) and hematoxylin. Then the slides were observed under the OLYMPUS BX53 Microscope and photographed.

### Statistical analysis

The statistical significance of differences between experimental groups was determined with two-tailed Student t test and analysis of variance using Prism 7.0 (GraphPad, San Diego, CA, USA). (*, P < 0.05; **, P < 0.01; ***, P < 0.001; ****, P < 0.0001).

## Acknowledgments

Thanks to Animal Center of Shanghai Jiao Tong University for providing the experimental platform. Thanks to Ms. Li Wei from the Public Experiment Center, school of pharmacy, Shanghai Jiao Tong University for her technical support. This research did not receive any specific grant from funding agencies in the public, commercial, or not-for-profit sectors.

## Authors’ Contributions

**W. Shi:** Conceptualization, methodology, investigation, data curation, formal analysis, writing-original draft. **L. Lv:** Validation, investigation. **N. Liu:** Resources, methodology, discussion. **H. Wang:** Supervision, investigation. **Y. Wang:** Investigation, formal analysis. **W. Zhu:** Validation, supervision. **Z. Liu:** Statistical analysis **J. Zhu:** Resources. **H. Lu:** Conceptualization, supervision, funding acquisition, writing-review & editing. All the authors read and approved the final manuscript.

